# Stable centrosomal roots disentangle to allow interphase centriole independence

**DOI:** 10.1101/199166

**Authors:** Robert Mahen

## Abstract

The centrosome is a non-membrane bound cellular compartment consisting of two centrioles surrounded by a protein coat termed the pericentriolar material (PCM). Centrioles must remain physically associated together (a phenomenon called centrosome cohesion) for cell migration, ciliary function and mitosis, yet how this occurs in the absence of a bounding lipid membrane is unclear. One model posits that pericentriolar fibres formed from rootletin protein directly link centrioles, yet little is known about the structure, biophysical properties or assembly kinetics of such fibres. Here, I combine live cell imaging of endogenously tagged rootletin with cell fusion, and find previously unrecognised plasticity in centrosome cohesion. Rootletin forms large, diffusionally stable, bifurcating fibres, which amass slowly on mature centrioles over many hours from anaphase. Nascent centrioles (procentrioles) in contrast do not form roots, and must be licensed to do so through polo-like kinase 1 (PLK1) activity. Transient separation of roots accompanies centriolar repositioning during the interphase, suggesting that centrioles organize as independent units, each containing a discrete root. Indeed, forced induction of duplicate centriole pairs allows independent re-shuffling of individual centrioles between the pairs. Thus, collectively, these findings suggest that progressively nucleated polymers mediate the dynamic association of centrioles as either one or two interphase centrosomes, with implications for our understanding of how non-membrane bound organelles self-organise.

## Introduction

The centrosome is a major microtubule organising centre, with critical roles in cell migration, cell division and cilia function. Mammalian interphase cells in G1 are generally thought to have one centrosome, consisting of two microtubule based structures, called centrioles. Centriole pairs are proteomically and functionally distinct [1], yet apparently remain physically associated, a phenomenon called centrosome cohesion [2–7]. Experimental changes to inter-centriolar distance during interphase result in defects in cell migration, ciliary function, and mitosis [8–12], underscoring the functional importance of centrosome cohesion. How two centrioles co-ordinately assemble into a single centrosome yet maintain distinct functions is largely unexplored.

Centrioles are not bounded by a lipid membrane but instead by two distinct structures, termed the pericentriolar material (PCM) and pericentriolar fibres [2,13]. Current models of PCM assembly emphasise high dynamics of constituent proteins, potentially as a liquid-like, toroidal structure [14,15]. In contrast, comparatively little is known about either the structure or assembly of pericentriolar fibres. Rootletin / ciliary rootlet coiled coil protein (gene symbol *CROCC*) localises to pericentriolar filaments, and rootletin knockout or knockdown results in both loss of filaments and centrosome cohesion [2,8,16,17]. One model posits that rootletin pericentriolar fibres directly connect centriole pairs to keep them spatially restricted [2,5,16,18]. Consistent with this proposal, rootletin is not found on mitotic centrosomes [5,18–20]. The kinetics of pericentriolar fibre dissolution during mitosis, when they reform, and the principles governing their replication are poorly understood, however.

To address these questions, this study uses high resolution imaging, genome editing and cell fusion to obtain unprecedented spatio-temporal information about the morphology, dynamics and assembly properties of rootletin fibres, referred to as roots. Roots are bifurcating adhesive structures which are licensed to form on centrioles by polo-like kinase 1 (PLK1) enzymatic activity. Both mature centrioles form independent roots which dynamically disentangle in response to organelle movement *in vivo*. Thus, they adopt a structure and function which allows centriole pairs to independently position during interphase, providing new insight into centrosome self-organisation.

## Results

### Centrosomal roots are large bifurcating fibres licensed to form on procentrioles by PLK1 activity

Pericentriolar filaments near centrosomes have been described for many decades [21], but their ubiquity in different cell types is unknown. To address this, the prevalence of rootletin fibres was documented by immunofluorescent staining and high resolution enhanced confocal Airyscan imaging in a range of cell types, whether cancerous, immortalised or primary. Thorough antibody validation, obtained by multiple independent lines of evidence, ensured specific recognition of rootletin (**Fig S1** and summarised in **Materials and methods**). Rootletin almost ubiquitously formed bifurcating fibres at the centrosome, henceforth referred to as roots (**Fig 1A**). Co-staining and segmentation of a range of markers of either centrioles or the PCM showed limited overlap with roots (**Fig 1B**), indicating they occupy a different locale, adjacent to the PCM and centrioles. Segmentation of both roots and centrioles, as marked by stable GFP-Centrin1 expression, showed that roots are large, at approximately ten-fold the size of a centriole on average in RPE cells (**Fig 1C**).

**Fig 1.**
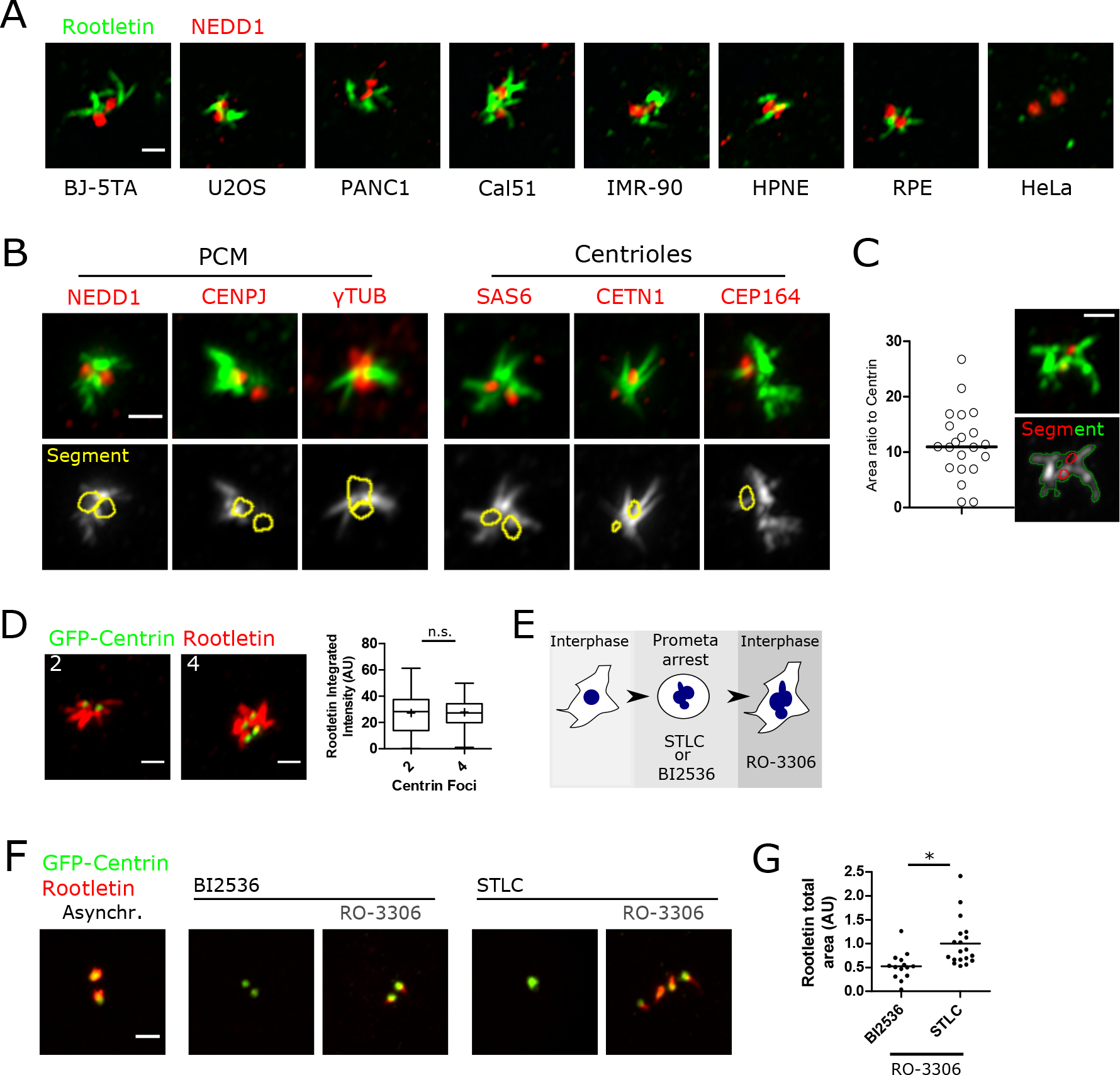
Centrosomal roots are large bifurcating fibres licensed to form on centrioles by PLK1 activity. (**A**) Anti-rootletin staining was imaged systematically in different cell types by Airyscan imaging (green). Pericentriolar material (PCM) is costained with anti-NEDD1 (red). Staining and imaging conditions are the same throughout. Confocal slices are shown. Scale bar 1μm. (**B**) Pairwise co-staining of rootletin (green) and other centrosomal genes (red), which are either in the PCM or centrioles as indicated. Maximum intensity projections, scale bar 1 μm. (**C**) Quantification of the ratio of rootletin immunostaining area relative to GFP-Centrin1 area from maximum intensity projected Airyscan images. (**D**) Rootletin immunofluorescent staining is equal in unreplicated centrosomes and diplosomes. Centrosomes were classified based on GFP-Centrin1 foci number (either two or four) and anti-rootletin staining was segmented. Scale bar 1μm. The mean is shown as + and the median as a horizontal bar. n.s., t-test. N=21 cells. Note that roots are shown in red in this panel. (**E**) Cells were arrested in prometaphase with either STLC (Eg5 inhibition) or BI2536 (PLK1 kinase inhibition), before being forced into interphase by RO-3306 (CDK1 inhibition), without the completion of mitosis. (**F**) Cells expressing GFP-centrin1 (green) were treated as depicted in (**E**) before staining with anti-rootletin antibody (red). Rootletin staining was not detected on prometaphase arrested cells. Maximum intensity projections are shown. Scale bar 1μm. (**G**) Root area per cell was quantified by direct segmentation of rootletin staining from images obtained as described in (**F**). * p=0.0006, t-test.

Centriole replication normally proceeds through the appearance of a nascent procentriole from the base of an existing centriole during S/G2 phase [22,23]. To examine whether procentriole formation influences root structure, interphase centrosomes were classified according to the presence of either two or four GFP-Centrin1 marked centrioles, corresponding to either unreplicated centrioles or diplosomes (mature centriole + nascent procentriole) respectively. No difference in rootletin intensity or size was detected (**Fig 1D**), suggesting that procentriole growth does not influence root structure.

Procentrioles mature into centrioles during mitosis, dependent on PLK1 activity, becoming replication competent after physically moving away from a centriole (a process termed disengagement) [24]. Therefore, the effects of PLK1 kinase inhibition on root formation were investigated, using the PLK1 kinase inhibitor BI2536 (**Fig 1E**). Cells arrested in mitosis through PLK1 blockade contained monopolar spindles [25], which were devoid of roots (**Fig 1F**; BI2536), consistent with previous work suggesting that mitotic cells do not have roots [5,16,18–20]. Since the inhibition of PLK1 results in cell cycle arrest, mitotic exit was forced into an ensuing interphase without cell division, by addition of the CDK1 inhibitor RO-3306 [26], to understand subsequent effects on root structure in interphase (**Fig 1E**). Control cells were also arrested in mitosis, but instead using the Eg5 kinesin motor inhibitor STLC followed by RO-3306. Control cells forced into interphase in this manner reformed roots despite unsuccessful mitotic genome segregation (**Fig 1F**; right hand panel entitled RO-3306). Interestingly however, forced mitotic exit after PLK1 blockade resulted in partial root reformation relative to STLC control (**Fig 1G**). These results suggest that centrioles are capable of root reformation in G1 regardless of PLK1 activity in the previous mitosis. In contrast, procentrioles must be modified by PLK1 dependent processes before they are competent to form roots in the next cell cycle. Furthermore, since PLK1 promotes centrosomal PCM expansion during mitosis [14], mitotic centrosomes disassemble roots even in the absence of centrosome maturation. Taken together, these results demonstrate that roots are large bifurcating fibres, that are found commonly in a range of cell types on mature PLKl-modified centrioles during the interphase.

### Diffusionally stable roots are progressively formed from anaphase

The dynamics and biophysical properties of eGFP tagged rootletin were investigated in living cells, by utilising both cDNA transgene overexpression and tagging of endogenous alleles. Consistent with previous work [2,16], over expression of eGFP-rootletin resulted in fibres and bifurcating fork structures which were longer than endogenous rootletin (e.g. compare **Fig S2A** with **Fig 1**). Time-lapse imaging of eGFP-rootletin fibre formation following transfection showed that eGFP-rootletin first appeared focally in a single location, prior to the emergence of a larger network over many hours (**Fig 2A**). Fibres increased in size not only by extension in length outwards from a single point, but additionally by the coalescence together of multiple fibres to form larger aggregates, frequently through end-on fusions (compare arrows in **Fig 2A**; **Video S1**).

**Fig 2.**
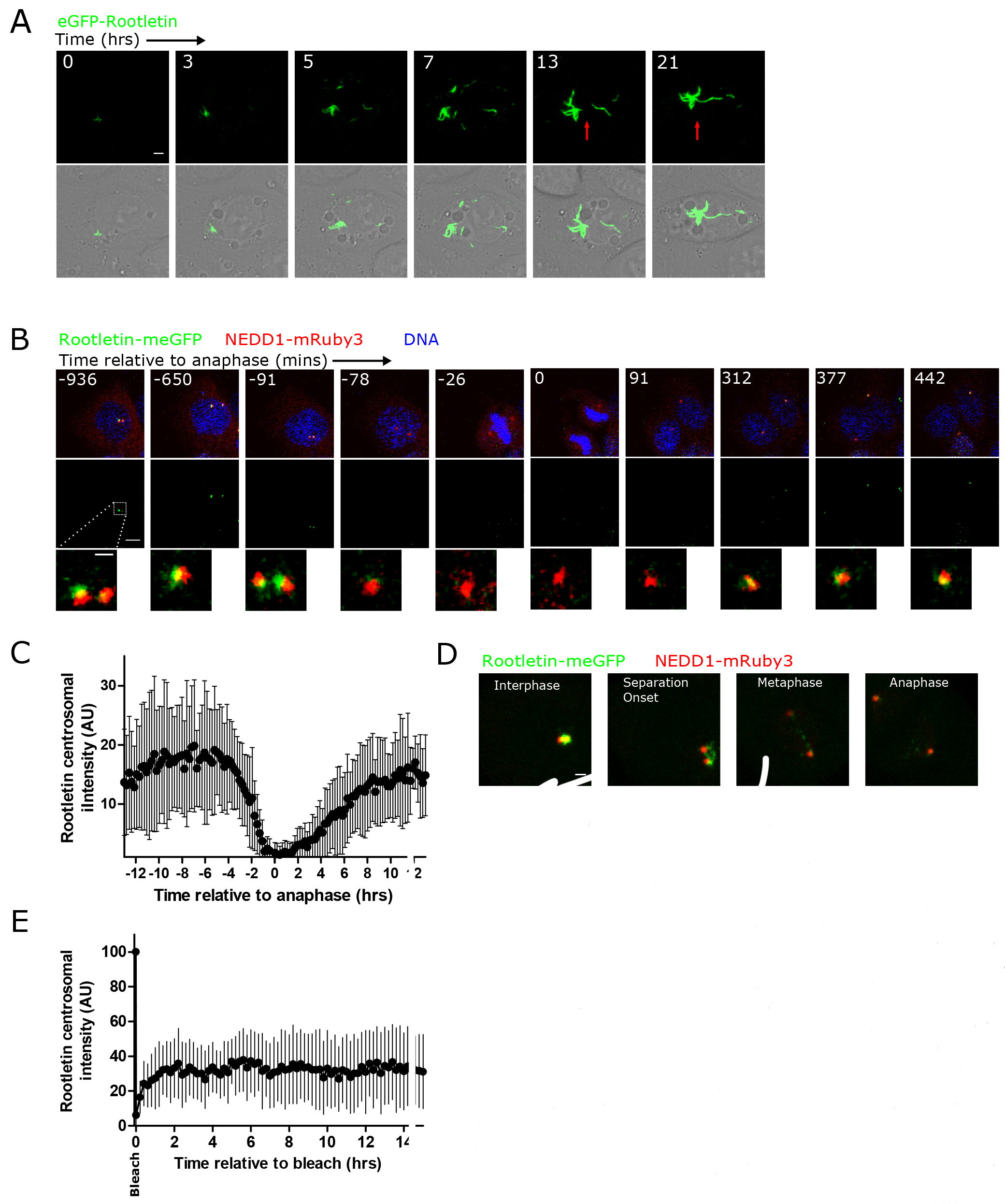
Diffusionally stable roots are progressively formed from anaphase. (**A**) eGFP-rootletin fibres progressively assemble following transfection. The images are timepoints from a single cell, taken by live cell 3D confocal time-lapse imaging. The arrows point to a fusion event of two pre-existing fibres. Scale bar 3μm. See also **Video S1** for the full timecourse. (**B**) Representative images from single cell three colour 3D confocal time-lapse imaging of rootletin-meGFP (green), NEDD1-mRuby3 (red; marking the PCM) and DNA (blue; marked by SiR-hoechst). Rootletin-meGFP (green) is visible at the centrosome during interphase but not during mitosis. Images were smoothed for display purposes here using a two-pixel median filter, but not for analysis. Scale bar 1 μm. See also **Video S2**. (**C**) Cell cycle dependent changes in rootletin-meGFP centrosomal fluorescence intensity. Centrosomes were automatically tracked based on NEDD1-mRuby3 as described in methods, to obtain the intensity of rootletin-meGFP in individual cycling cells. Traces were manually aligned relative to anaphase onset based on SiR-hoechst staining of DNA (time 0) to create a plot of the mean +/− SD, N=17 cells. (**D**) An example of rootletin-meGFP (green) during centrosome separation in early mitosis. NEDD1-mRuby3 is shown in red as centrosomes move apart. Scale bar 2μm. (**E**) FRAP recovery curve over 15 hours, plotting the mean +− SD centrosomal intensity of rootletin-meGFP from 3D confocal imaging after fluorescence bleaching, in thymidine arrested cells. Centrosome position was efficiently tracked independently of rootletin-meGFP fluorescence intensity through the use of simultaneous NEDD1-mRuby3 imaging in a spectrally distinct channel. N=11 cells.

Cell cycle dependent changes in the centrosomal intensity of meGFP tagged rootletin were followed by 3D confocal time-lapse imaging. Since overexpressed rootletin fibres were larger than endogenous antibody stained roots, and because overexpression can influence quantitative measures of protein function *in vivo* [27], CRISPR Cas9 was used to insert an in-frame fusion of meGFP into the endogenous rootletin (*CROCC*) locus and therefore study rootletin behaviour with live cell microscopy at endogenous levels for the first time (**Fig S3**). Homozygous tagging in the diploid breast cancer cell line Cal51 resulted in fluorescent signal closely resembling antibody staining (**Fig S3E**). Rootletin-meGFP was barely detectable at the centrosome during mitosis, consistent with immunofluorescent staining (**Fig 1**), and consequently, stable co-expression of a NEDD1-mRuby3 fluorescent fusion marking the PCM was used to track centrosomes throughout the cell cycle and independently of rootletin levels, in a spectrally distinct fluorescent channel (**Fig 2B**; **Video S2**). Additionally, fluorescently labelled chromatin was imaged, to reveal mitotic sub stage.

Rootletin began to be released from the centrosome more than two hours prior to anaphase (**Fig 2C**). By anaphase, centrosomal rootletin could not be detected above cytoplasmic levels, suggesting disassembly of all centrosomal roots. Rootletin centrosomal levels increased from anaphase, but unexpectedly, continued to increase at a slow rate for approximately nine hours and thus significantly into G1 phase. Staging of rootletin intensity relative to centrosome separation revealed that its release from the centrosome began prior to centrosome separation and continued after it, with low levels of rootletin still present during centrosome separation which could be ripped apart during poleward centrosome migration (**Fig 2D**). These results suggest that the removal of rootletin from centrosomes begins early in mitosis or in late G2 phase, prior to both chromatin condensation and centrosome separation, and then continues during these processes. Rootletin assembly at the centrosome begins from anaphase and occurs slowly for approximately nine hours into G1 phase.

Some centrosomal PCM components show dynamic exchange of subunits on the seconds timescale - a property which has been suggested to be important for centrosome assembly [13]. Fluorescence recovery after photobleaching (FRAP) was therefore used to ask whether rootletin forms steady state polymers. FRAP of extended eGFP-rootletin fibres showed almost no movement of eGFP-rootletin over a time period of ten minutes however, even after a relatively rapid bleach (**Fig S2B**; ~1 sec). Lack of recovery was not due to image bleaching or fibre movement out of the field of view, since adjacent unbleached ends of the fibre remained unchanged. To investigate very slow dynamic exchange of endogenous centrosomal rootletin-meGFP, on the hours timescale, cells were arrested at G1/S of the cell cycle using thymidine, to circumvent the effects of cell cycle progression on root morphology (see **Fig 2C**). Centrosomal rootletin-meGFP fluorescent signal was then bleached, and recovery followed by tracking of NEDD1-mRuby3 marked centrosomes during time-lapse imaging (**Fig 2E**). Recovery of rootletin-meGFP fluorescence on this long timescale was limited to ~30%. Therefore, eGFP-rootletin fibres are predominantly diffusionally stable structures which are slowly and progressively assembled over hours following anaphase.

### Roots disentangle during transient centriole splitting in interphase

How a single interphase cell co-ordinately organises two disengaged centrioles is unclear. The prevalence of centrosomal cohesion was systematically documented in a range of human tissue culture cell types by automated fluorescence imaging and analysis (**Fig 3A**). Quantification of the percentage of cells with split centrosomes - defined as two PCM foci >1.5 μm apart - showed it was low at ~10%, dependent on cell type (see **Materials and methods** for a definition of split centrosome). Thus, in most cell types centrioles remain cohered in close proximity during interphase, consistent with previous work [6,28–32]. One possibility is that the minority of cells with split centrioles keep them stably separated over time, perhaps due to a permanent failure of centrosome cohesion. Interestingly however, single cell 3D confocal live imaging of centriole pairs marked by GFP-Centrin1 showed transient splitting. Hence, a single centriole pair would split into two and then rejoin, often repeatedly (**Fig 3B**; **Video S3**). Transient centriole splitting was manifest in live cell imaging of several different cell types including Cal51, HeLa, RPE and U2OS cells (**Fig 3B-3E**; **Video S4** and **S5**).

**Fig 3.**
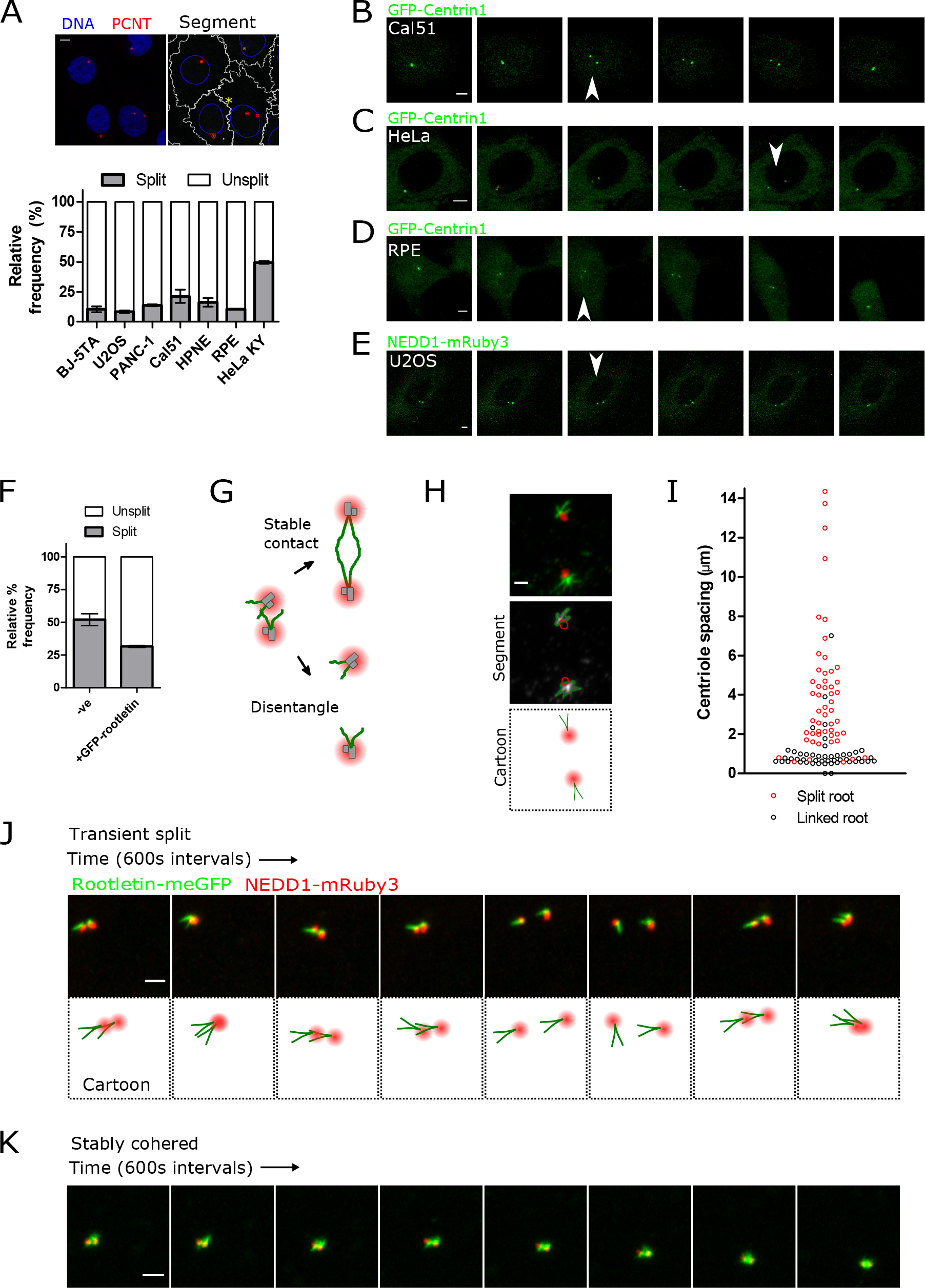
Roots disentangle during transient centriole splitting in interphase. (**A**) Quantification of centrosome cohesion in the interphase of various cell types through systematic immunofluorescent staining. The images show representative staining of PCNT (red; marking centrosomal PCM) and DNA (blue; hoechst 44432). Scale bar 5μm. The right panel shows representative segmentation of centrosomes (red), nuclei (blue) and cytoplasm (white) in Cal51 cells. The yellow asterisk denotes a cell containing two centrosome foci. The bar graph shows the mean % of cells with centrosomes separated by >1.5μm, from a minimum of 500 cells of each cell type. Error bars show SEM from two experiments. (**B - E**) Selected frames showing centriole splitting in live 3D confocal time-lapse imaging. Centrosomes are marked by either GFP-Centrin1 or NEDD1-mRuby3. Arrows denote centriole splitting events. The time intervals between frames are 12 minutes (**B, C**), 24 minutes (**D**) or 8 minutes (**E**). Scale bar 5μm. See also **Videos S3-S5**. (**F**) Centrosome cohesion in HeLa cells -/+ overexpression of eGFP-rootletin, measured by automated imaging and analysis. Horizontal bars show the mean of two experiments +− SEM. * p<0.001 by Fischer’s exact test. More than 1000 cells were measured for each sample. (**G**) Opposing models of root behaviour during centriole splitting, termed “Stable contact” or “Disentangle”. (**H**) Representative image of root disentanglement after centriole splitting. Scale bar 1μm. (**I**) Root linkage plotted as a function of centriole spacing distance. (**J, K**) High resolution Airyscan time-lapse imaging of endogenous rootletin-meGFP and NEDD1-mRuby3 during a centriole split (**J**) and when remaining stably cohered (**K**) in Cal51 cells. Scale bar 2μm. See also **Videos S6** and **S7**.

In agreement with a published report [29], HeLa Kyoto cells had high levels of centrosome separation, with ~50% of cells showing split centrioles in a fixed asynchronous population (**Fig 3A**). Since HeLa cells have low levels of rootletin expression and short roots (**Fig 1A**), and previous work has shown that rootletin knockdown results in the loss of centrosome cohesion [2,33], the effect of increasing root length on centrosome position was investigated in HeLa cells, to ask whether rootletin over expression is sufficient to increase centrosome cohesion. An increase in rootletin fibre length significantly increased centrosome cohesion in interphase HeLa cells, as measured by automated imaging and analysis of immunofluorescently stained samples (**Fig 3F**, p<0.001, Fischer’s exact test). Together these results show that although centrioles generally remain cohered into a single focal location, they are able to transiently split apart in interphase in a manner that is antagonised by eGFP-rootletin over expression.

How might rootletin fibres respond to transient centriole splitting? Two opposing models for root behaviour after centriole splitting were postulated (**Fig 3G**). The first is maintenance of a stable root contact between centrioles as they move apart, for example due to stretching. The second is loss of physical connection and disentanglement (“Stable contact” versus “Disentangle” respectively). Surprisingly, and in contrast to cohered centrosomes, split centrioles were not linked by rootletin fibres (**Fig 3H**). Instead, a transition from linked to unlinked roots occurred at distance greater than ~1.5μm between centrioles (**Fig 3I**), thus supporting the disentanglement model.

Simultaneous two-colour Airyscan microscopy of root disentanglement in living cells revealed that roots occupy markedly heterogenous orientations which change in response to *in vivo* centriole movement (**Fig 3J**; **Video S6**). The centrosome distal ends of roots have the capacity to pivot relative to centrosome proximal ends, suggesting a common more stable attachment point at the proximal end. Pivoting of centrosome distal tips was not just observed in centrosomes with split centrioles but also in cohered centrosomes, with roots maintained stably at the centriole-centriole interface (**Fig 3K**; **Video S7**). As centrosomes remerged after a split, roots did not necessarily join, but could alternatively contact the PCM of the opposing centriole. Together these observations indicate that although roots can be maintained stably at the interface between centrioles, their orientation is heterogeneous, and notably, in response to centriole movement they disentangle rather than maintaining a continuous direct linkage.

### Centriole independence during interphase

Since disengaged centrioles can transiently split (**Fig 3**), the comparative structure of roots and PCM on centrioles was investigated in detail during splitting. Root area was approximately halved in split versus cohered centrioles (**Fig 4A**), suggestive of equal partitioning of two independent roots. Due to their mode of replication, mature centrioles are of differing age, with the oldest centriole marked by appendage proteins including CEP164 [34]. Discrimination of centriole age using CEP164 immuno-staining showed that roots are nucleated symmetrically on both centrioles when spatially separated (**Fig 4B**). A similar comparison of PCM structure with the PCM resident pericentrin showed similarly that centrioles individually nucleate PCM when split (**Fig 4C**), something also evident in live cell imaging of NEDD1-mRuby3 (**Fig 3J-K**), and previous work [24,30,32]. These observations show that centrioles independently maintain roots and PCM during centrosome splitting in interphase.

**Fig 4.**
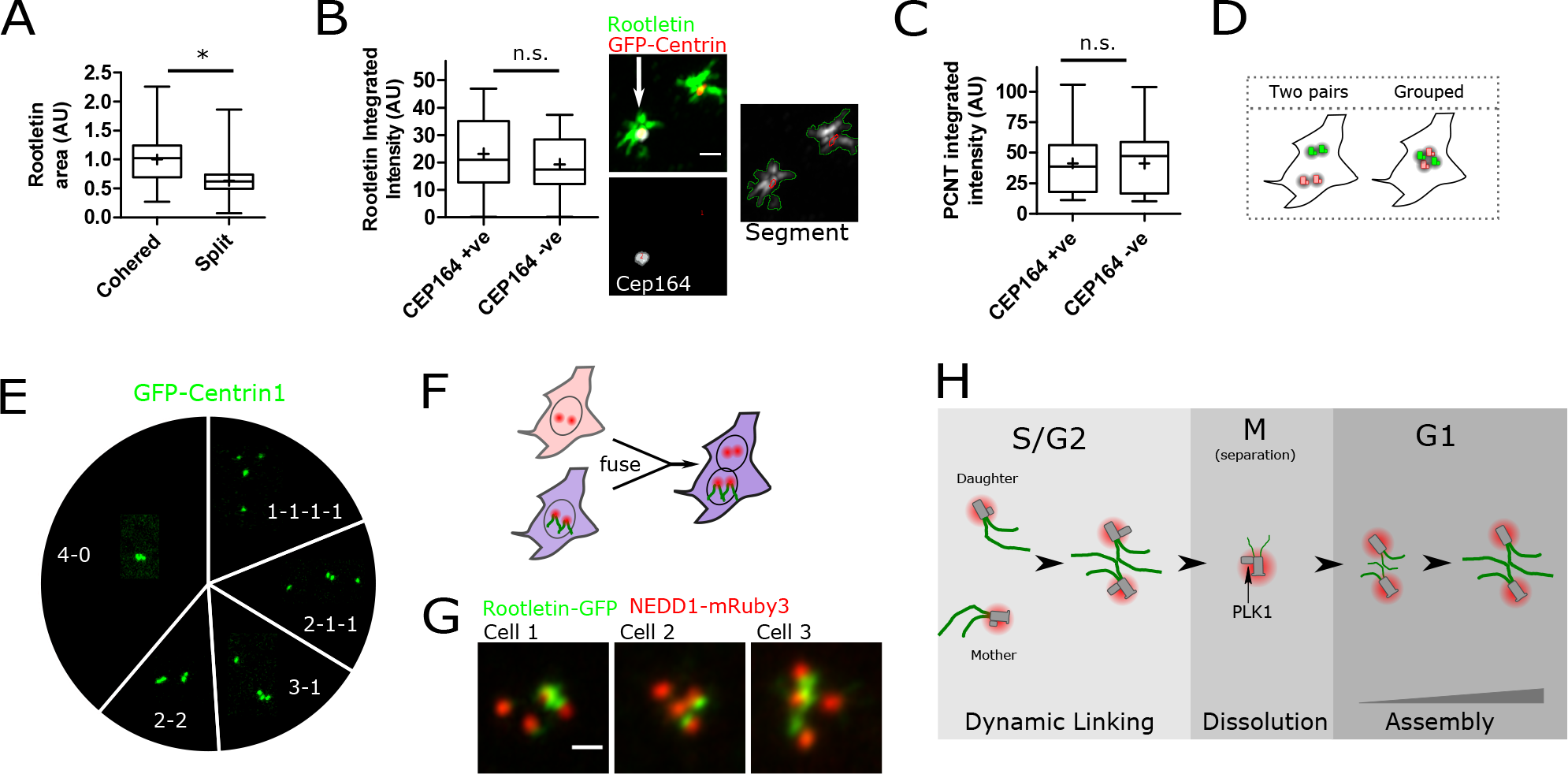
Centriole independence during interphase. (**A**) Root fibre area is significantly lower in split versus cohered centrioles (p<0.0001, t-test). Anti-rootletin immunofluorescent staining was Airyscan imaged and segmented, N=36 cells from two experiments. (**B**) Rootletin immunofluorescent staining (green) is the same at both mature centrioles (n.s., t-test). Centrioles were identified based on GFP-Centrin fluorescence (red), and then classified according to age, identifying the older centriole by CEP164 positivity (white). The arrow denotes a CEP164 positive centriole. N=21 cells per sample. Scale bar lμm. (**C**) PCNT immunofluorescent staining (of the PCM) is the same on either mature centriole (n.s., t-test). Cells were imaged and analysed as described in **B**, except segmenting PCNT staining. N=21 cells. (**D**) Cells with four mature centrioles might either maintain them as separate pairs or cohere them together. (**E**) The pie chart shows the proportion of each GFP-Centrin1 centriole configuration in cells with four centrioles, produced as depicted in **Fig 1E** - by sequential arrest in mitosis by STLC treatment, followed by induction into interphase with RO-3306. The images are representative of each configuration and the text denotes the configuration, e.g. 40 indicates all four centrioles cohered in one location. (**F**) Cells expressing rootletin-meGFP were stained with CellTrace Violet dye and then fused with cells stably expressing NEDD1-mRuby3. (**G**) Root arrangement in three different fused cells, produced as described in (**F**). Note that four NEDD1-mRuby3 foci are visible due to dynamic exchange of NEDD1 between the centrosome and cytosol. Scale bar lμm. (**H**) Interphase centriole pairs contain large bifurcating fibres which disentangle when centrioles move apart >1.5μm relative to each other. Root dissolution begins prior to mitotic centrosome separation and chromosome condensation. At the time of centrosome separation, roots are diminished in quantity and ripped apart during poleward movement of centrosomes. Roots form slowly over many hours from anaphase, as diffusionally stable fibres. PLK1 dependent modification of procentrioles allows root formation on mature centrioles in the ensuing cell cycle.

Given the dynamic nature of centrosome cohesion (**Fig 3**) and root disentanglement, it was of interest to investigate whether mature centriole pairs would be maintained in cells with four mature centrioles (**Fig 4D**). Interestingly, centriole position in cells forced into interphase after a failed mitosis by STLC treatment (**Fig 1E**), showed all possible centrosome cohesion configurations. Most commonly, all four centrioles grouped together as one (**Fig 4E**; ~40%), but notably other spatial arrangements were equally as likely as two pairs. Thus, centrioles are not maintained separately in stable pairs but will cohere together, even in a grouping such as one single centriole and three cohered separately (labelled as “3-1” on **Fig 4E**). This was examined further using polyethylene glycol mediated cell fusion of two different cell types, one expressing rootletin-meGFP and the other not (see **Materials and methods** for details). Fused cells contained two nuclei and four mature centrioles - one mature centriole pair from each different cell line (**Fig 4F**). As per after mitotic failure (**Fig 4E**), after cell fusion centriole pairs were not exclusively maintained, despite their origin in different cells (**Fig 4G**). Instead, fluorescent roots frequently embraced all four centrioles. Therefore, by two independent methods, mature centriole pairs are not stably maintained in cells with four centrioles.

## Discussion

Cells must carefully regulate centrosome number and position, coordinating two centrioles which are capable of distinct functions [23,30,35]. The data here provoke an interesting hypothesis; that interphase cells always have two centrosomes which are generally held together by stable fibres which reach outward into the cytoplasm. Three key pieces of evidence support this model. Firstly, both mature interphase centrioles in a pair independently nucleate roots, as well as PCM. Secondly, these units - consisting of a centriole - root - PCM, have the capability to transiently spatially separate during interphase, a phenomenon accompanied by root disentanglement. Thirdly, cells engineered with two centriole pairs do not maintain them separately, but instead dynamically make new groupings. Thus, there is remarkable plasticity in the maintenance of centrosome cohesion, with individual centrioles able to rearrange between pairs, through dynamic splitting of roots. These conclusions are consistent with previous observations of split centrioles [6,28–32]. Centriole independence could aid plasticity of centrosome function such that the two centrioles can either act as one or separately. Thus, the data presented here explains previous observations that centrioles may have either different or coordinated functions [23,30,35].

How non-membrane bound organelles regulate their position, size and number within the cellular interior is a major open question. Recent work has postulated that organelles such as centrosomes phase separate as liquid-like compartments [36]. This model is characterised by high internal turnover of components parts, spherical shape, and the ability of multiple organelles to fuse [15]. In contrast, roots are diffusionally stable, remain separate through multiple cycles of merging and splitting, and are not spherical, instead potentially engendering polarity to the centrosome as a branched organelle. Hence, roots have surprisingly different organisational principles to the PCM. Further work will be needed to understand whether this has implications for how centrosomal position is regulated; a testable prediction from this work is that the two PCM clouds could merge with liquid like properties as centrosomes cohere.

In conclusion, root mediated splitting of two centrosomes might allow plasticity of cytoskeletal function, thus explaining how two non-membrane bound organelles co-ordinately function in either one or two locations during the interphase [28,32,37]. It is tempting to suggest that progressively nucleated, diffusionally stable polymers might be used generally to maintain organelle subcellular position and number [38].

## Materials and methods

### Antibody validation

Multiple lines of evidence indicate that a commercially available anti-rootletin antibody (Novus Biologicals NBP1-80820) specifically recognises the product of the *CROCC* gene. siRNA depletion of rootletin (*CROCC*) using RNA interference removed signal by both immunofluorescence and western blot in multiple cell types (**Figs S1A**, **S1B** and **S1D**). An antibody independent method, GFP tagging, showed significant overlap to protein abundances measured by immunofluorescence, both in time and space (this is apparent throughout Figs 1**-3**). For example, rootletin signal was virtually undetectable in metaphase by either antibody or GFP tagging. Finally, anti-rootletin antibody also stained GFP-rootletin when overexpressed as a transgene (**Fig S1C**).

### Immunofluorescence

Cells were fixed in 4% paraformaldehyde or ice cold 100% methanol for ten minutes, permeabilised in 0.1% triton and blocked in 3% bovine serum albumen (Thermo Fisher Scientific). Antibodies used were: rabbit anti-CROCC (1:250-1:750, Novus Biologicals NBP1-80820), mouse anti-NEDD1 (1:500, Abcam ab57336), rabbit anti-PCNT (1:1000; Abcam ab4448), mouse anti-SAS6 (1:300, Santa Cruz Biotechnology sc-81431), mouse anti-CENPJ (1:100, Santa Cruz Biotechnology sc-81432), mouse anti-gamma Tubulin (1:1000, GTU-88, ice cold methanol fixation), mouse anti-CETN1 (1:4000, EMD Millipore 20H5), mouse anti-CEP164 (1:200, Santa Cruz Biotechnology sc-515403).

### Cell culture, chemicals and DNA constructs

Cal51 (German Collection of Microorganisms and Cell Cultures ACC303), U2OS (American Type Culture Collection ATCC HTB-96), HeLa Kyoto, PANC-1, and IMR-90 cell lines were grown in Dulbecco's modified Eagle's Medium (DMEM) supplemented with 10% fetal calf serum, GlutaMAX (Thermo Fisher Scientific), 100ug/ml penicillin/streptomycin. hTERT RPE1 cells were cultured in DMEM/F12 with 10% FBS, Penicillin/Streptomycin and 4.2% sodium bicarbonate. h-TERT BJ-5ta (ATCC CRL-4001) were grown in a 4:1 mixture of DMEM to M199. h-TERT HPNE (ATCC CRL-4023) were grown in a 3:1 mixture of DMEM to M3:BaseFmedium, with 5% fcs, 10ng/ml EGF, 2mM glutamine and 750ng/ml puromycin. For live cell imaging cells were cultured in either phenol red free Fluorbrite DMEM medium (Thermo Fisher Scientific) or L15 CO2 independent medium, on Ibibtreat coated Ibidi μ-Slide 8-well dishes (Ibidi). All tissue culture reagents were purchased from Sigma-Aldrich unless stated otherwise.

Transfection was with lipofectamine 3000 (Invitrogen) according to the manufacturer's instructions. SiR-Hoechst (Tebu Bio) was used at 200nM and incubated for 30 minutes at culture conditions before replacing with fresh medium for imaging. siRNA knockdown of rootletin (*CROCC*) was with Dharmacon ON-TARGETplus SMARTpool, transfected with RNAiMax transfection reagent (Thermo Fisher Scientific). NEDD1-mRuby3 was synthesised by Thermo Fisher Scientific as codon optimised sequence in the vector pcDNA 3.1(+), and contained the reference sequence of human NEDD1, linked to fluorescent protein by five glycine residues.

### cDNA stable cell line production and CRISPR Cas9 mediated genome editing

Stable cell lines expressing cDNA constructs were produced by transfection followed by culture for > four days, either with or without antibiotic selection, followed by fluorescence-activated cell sorting (FACS) of fluorescent cells relative to matched untransfected negative controls. Genome editing was essentially as described in [27], with some modifications. Guide RNA directing CRISPR Cas9 mediated DNA damage was expressed from pX330-U6-Chimeric_BB-CBh-hSpCas9 (Addgene plasmid #42230). Guide RNA sequences all overlapped the *CROCC* STOP codon against the +ve strand they were as follows (5' ---- 3'): CCAGCAGGAGCTCATTTCTC, CCAGAGAAATGAGCTCCTGC, CAGGAGCTCATTTCTCTGGG. To insert meGFP into the *CROCC* gene locus, a donor plasmid was constructed in the vector pUC19 by HD In-fusion cloning (Clontech). It consisted of 700bp homology arms from the C-terminus of the *CROCC* genomic reference sequence, surrounding the meGFP coding sequence. Five glycine residues linked the C-terminus of the gene to the fluorescent protein. Insertion of meGFP into the endogenous *CROCC* locus was detected by extraction of genomic DNA using QuickExtract DNA extraction solution (epicentre) according to the manufacturer’s instructions, followed by junction PCR with the following primers; Forward: GGCTGGCCTTACCTTCCCTT, Reverse: CTGGAAGGCCTGTCACTGTC.

### Mitotic arrest and release

Cells were arrested for 12 hours in either 200nM BI2536 (Sigma-Aldrich) or 10μM S-trityl-L-cysteine (STLC). Only mitotically arrested cells were analysed further, by mitotic shake-off. Mitotic exit was then forced with R0-3306 (10μM) for six hours. Cells were confirmed as being in interphase during interphase.

### Image analysis

Images are presented as maximum intensity projections from 3D data unless otherwise stated. Image brightness and contrast settings were changed linearly and consistently between samples for display purposes of representative images, but not for quantitation.

The intensity of centrosomal rootletin-meGFP in cycling cells was determined by automated centrosome tracking after movie acquisition. Centrosomes were segmented and tracked using the Trackmate plugin in ImageJ / Fiji [39], using LAP Tracker, and confirmed as successful by manual analysis of tracking. NEDD1-mRuby3 was tracked, a marker of the PCM which was found to be present throughout the cell cycle. Individual cell tracks were aligned manually relative to anaphase, or the nearest frame to anaphase, based on both bright-field and SiR-hoechst fluorescent DNA labelling.

Segmentation from fixed images was in Cell Profiler software, with data analysis in Knime software, using custom built analysis pipelines. For calculation of per cell centriole splitting, nuclei were detected based on hoechst staining, and cytoplasm using a watershed algorithm outwards from nuclei based on gamma-tubulin staining. Mitotic cells were excluded on the basis of hoechst staining, since mitotic DNA was smaller and more densely stained than in interphase nuclei. Centrosomes were detected with PCNT staining and defined as split if a cell contained two PCNT foci >1.5μm apart by Euclidean straight-line distance. 1.5μm was chosen as the definition of split centrioles since this distance was the threshold above which roots rarely linked centrioles in high resolution imaging (**Fig 4D**).

For segmentation of roots, various thresholding strategies were used in CellProfiler, including propagation outwards from a GFP-centrin1 seed region, or direct thresholding. Spacing of PCM staining was measured by adaptive thresholding followed by calculation of 2D Euclidean distance between centroids. In this case roots were segmented using propagation from PCM and then manually classified as linked if one pixel overlap occurred between a root from each PCM.

### Western blotting

Cells were lysed for 20 minutes on ice in RIPA buffer (50 mM Tris HCl, pH 8, 150 mM NaCl, 1% NP40, 0.5 M sodium deoxycholate, 0.1% SDS, complete protease inhibitor cocktail, PhosSTOP (Roche)). Protein concentration was quantitated using the bicinchoninic acid method (Sigma-Aldrich). Whole cell extracts were separated by electrophoresis on a 3-8% Tris-Acetate gel and transferred to PVDF membrane using the iBlot system (Thermo Fisher Scientific) according to the manufacturer’s instructions. Membranes were blocked in 5% milk dissolved in 0.1%Tween/TBS. Antibodies used were rabbit anti-CROCC (1:250-1:750 overnight; Novus) and mouse monoclonal beta-Actin (1:10,000 one hour at room temperature; Sigma-Aldrich).

### Live cell time-lapse imaging and fluorescence recovery after photobleaching

Cells were imaged without phenol red in either L15 CO_2_-indepdendent medium, or in Fluorbrite Imaging medium with 5% CO_2_ at 37°C, in Ibidi u-slide 8-well dishes. Imaging was with a Carl Zeiss 880 Airyscan, either in Airyscan or standard confocal mode, using either a 63x NA 1.4 or 100x NA 1.4 oil immersion lens. FRAP was performed essentially as described in [14], by bleaching using a 488 argon laser at 100% for the minimum time required to cause ~50% fluorescence loss (but keeping the bleach time the same duration in all samples). Corrections were made for non-specific image bleaching by using time courses taken with identical settings but in unbleached cells. Hence, lack of recovery was not simply due to nonspecific bleaching. In long FRAP experiments (15 hours total recovery period), significant movement occurred, both of the centrosome and in some cases the whole cell. To account for this, centrosome position was tracked by using a spectrally distinct centrosomal marker, NEDD1-mRuby3.

### Cell fusion

Cal51 cells expressing rootletin-meGFP were fused with cells expressing NEDD1-mRuby3. Prior to fusion, rootletin-meGFP cells were stained with CellTrace Violet dye (Thermo Fisher Scientific), according to the manufacturer’s instructions. This improved the identification of fused cells by FACS as detailed below. Fusion was by mixing trypsinised cells at a 1:1 ratio and incubating for five minutes with Hybri-Max 50% 1450 polyethylene glycol solution (Merck). Serum free medium was then added dropwise for one minute before 10 minutes incubation at 37°C with normal medium. Fresh medium was then added before a two-hour incubation at 37°C. Fused cells were at low frequency (<1%), and so were enriched by FACS sorting, by gating for both CellTrace Violet (Thermo Fisher Scientific) and NEDD1-mRuby3 positivity relative to negative controls. Since neither cell line originally had both colours, this strategy allowed identification of fused cells. Fused cells were sorted by flow cytometry directly into an Ibidi μ-Slide Angiogenesis imaging dish and confirmed by microscopy as aneuploid relative to the single colour lines, as expected. Note that fused cells contained up to four centrioles marked by NEDD1-mRuby3 fluorescence, due to turnover of this marker at the centrosome from the cytoplasmic pool. Thus, upon fusion of two cells, NEDD1-mRuby3 was able to bind centrosomes through recruitment from the cytoplasmic pool. Since rootletin-meGFP shows very slow diffusional exchange (**Fig 2**), this process did not occur to the same extent.

## Supporting information

**Video S1.** Growth of cDNA eGFP-rootletin fibres in a single Cal51 cell after transfection. Each frame is taken at a six-minute interval and shows a maximum intensity z-projection from a 3D confocal stack.

**Video S2.** Cell cycle dependent changes in centrosomal rootletin-meGFP intensity (green; roots) in Cal51cells coexpressing NEDD1-mRuby3 (red; marking the PCM) and stained with SiR-Hoechst (blue; DNA). Each frame is taken at a 12-minute interval and shows a maximum intensity z-projection from a 3D confocal stack.

**Video S3.** Centriole splitting and cohesion visualised by 3D confocal time-lapse imaging of GFP-Centrin1 (centrioles) in Cal51 cells. Each frame is taken at a 12-minute interval and shows a maximum intensity z-projection. Note that this cell divides after 25 frames.

**Video S4.** Centriole splitting and cohesion visualised by 3D confocal time-lapse imaging of GFP-Centrin1 (centrioles) in HeLa cells. Each frame is taken at a 12-minute interval and shows a maximum intensity z-projection.

**Video S5.** Centriole splitting and cohesion, visualised by 3D confocal time-lapse imaging of GFP-Centrin1 (centrioles) in RPE cells. Each frame is taken at a 24-minute interval and shows a maximum intensity z-projection.

**Video S6.** Root disentanglement during centriole splitting and remerging, visualised by 3D confocal Airyscan time-lapse imaging of rootletin-meGFP (green; roots) and NEDD1-mRuby3 (red; PCM). Each frame is taken at a 10-minute interval and shows a maximum intensity z-projection.

**Video S7.** Root behaviour in a stably cohered centrosome, visualised by 3D confocal Airyscan time-lapse imaging of rootletin-meGFP (green; roots) and NEDD1-mRuby3 (red; PCM). Each frame is taken at a 10-minute interval and shows a maximum intensity z-projection.

**Fig S1.**
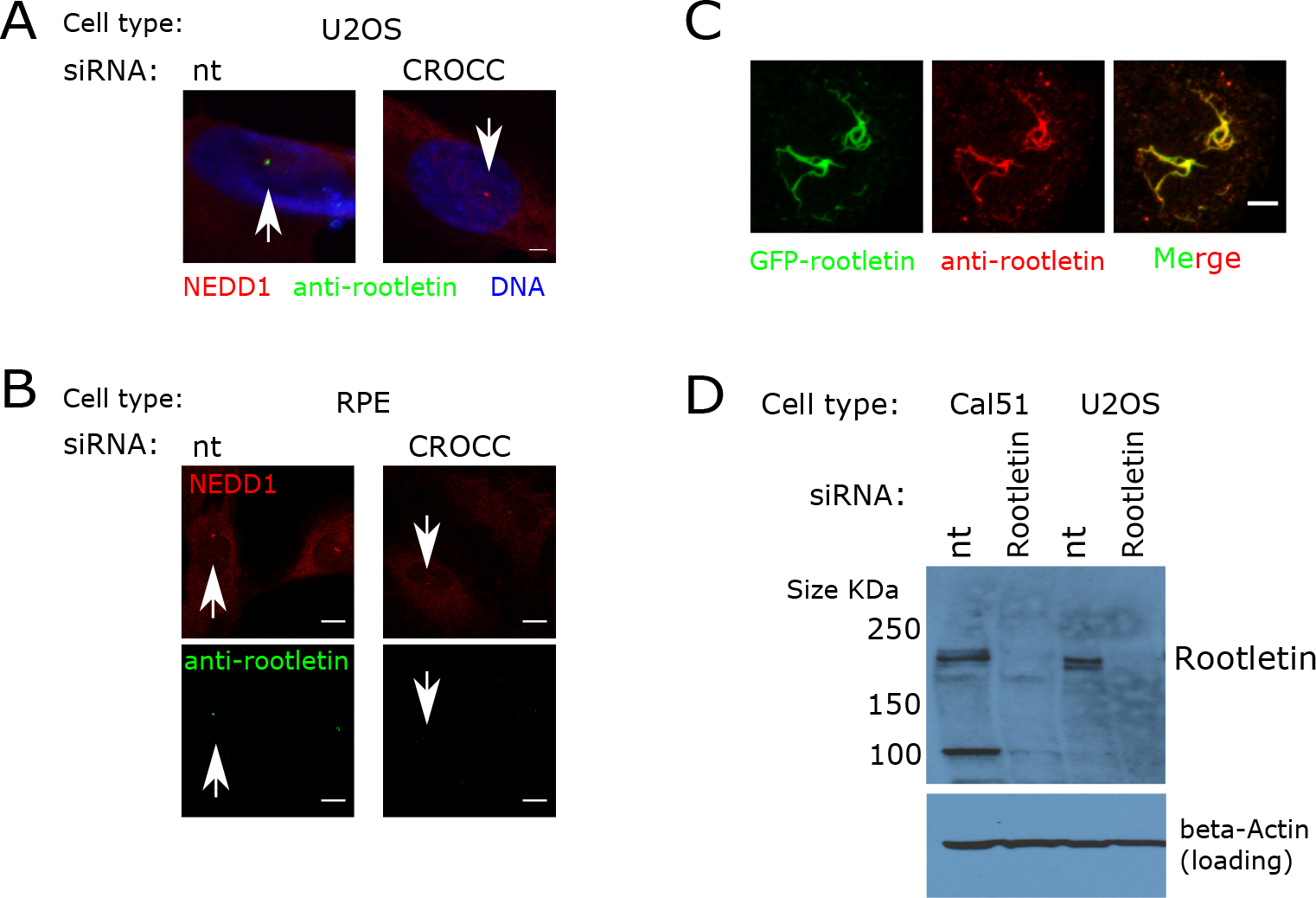
Validation of anti-rootletin antibody (related to Fig 1). (**A, B**) Anti-rootletin immunofluorescent staining (green) is not evident at centrosomes after rootletin (*CROCC*) siRNA. Centrosomes were co-stained with anti-NEDDl antibody (red), in multiple cell types. Anti-rootletin staining (green) is present after non-targeting siRNA negative control (nt) treatment. Arrows have been annotated manually to indicate centrosomes. Imaging conditions and brightness and contrast settings are consistent between control and siRNA treated samples. (**C**) Anti-rootletin antibody stains eGFP-rootletin over expressed from a cDNA transgene. Anti-rootletin staining is shown in red and GFP-rootletin fluorescence is shown in green. (**D**) Anti-rootletin bands are not detected by western blot of whole cell lysate after rootletin (*CROCC*) siRNA, demonstrating antibody specificity in multiple cell types.

**Fig S2.**
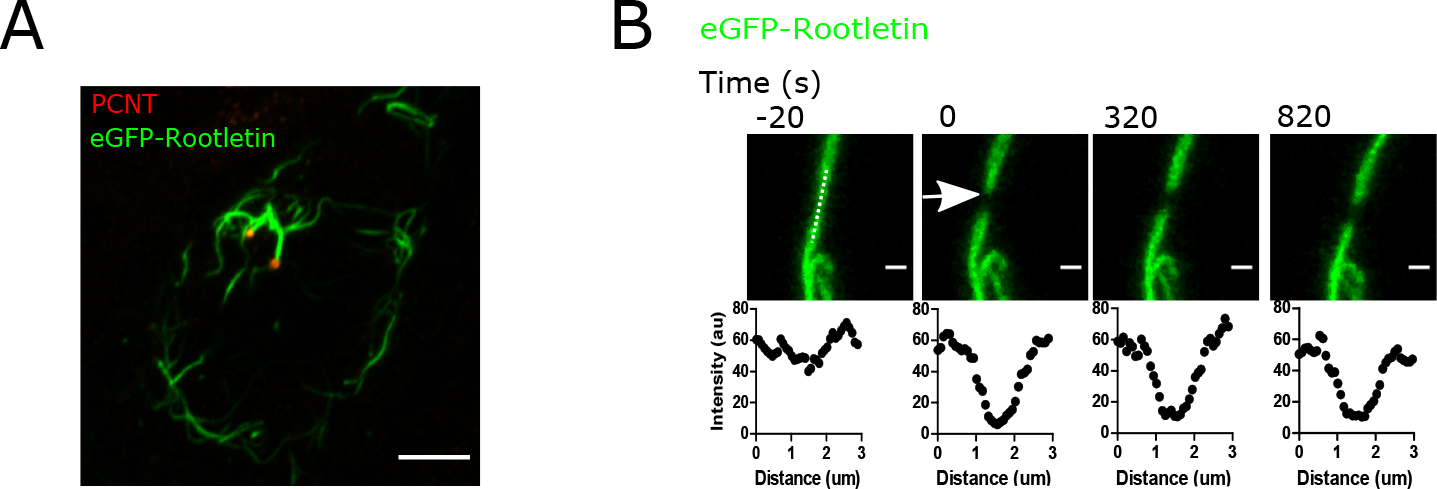
Over expression of GFP-rootletin progressively assembles large fibres which are diffusionally stable over minutes (related to Fig 2). (**A**) Representative maximum intensity z-projection Airyscan image of over-expressed eGFP-rootletin fibres (green), co-stained with the PCM marker PCNT (red). Scale bar 5μm. (**B**) FRAP of a single eGFP-rootletin fibre in the location denoted by the arrow. The graphs show the fluorescence intensity along a line profile at each timepoint, denoted by the dashed line in timepoint −20s. Scale bar 3μm.

**Fig S3.**
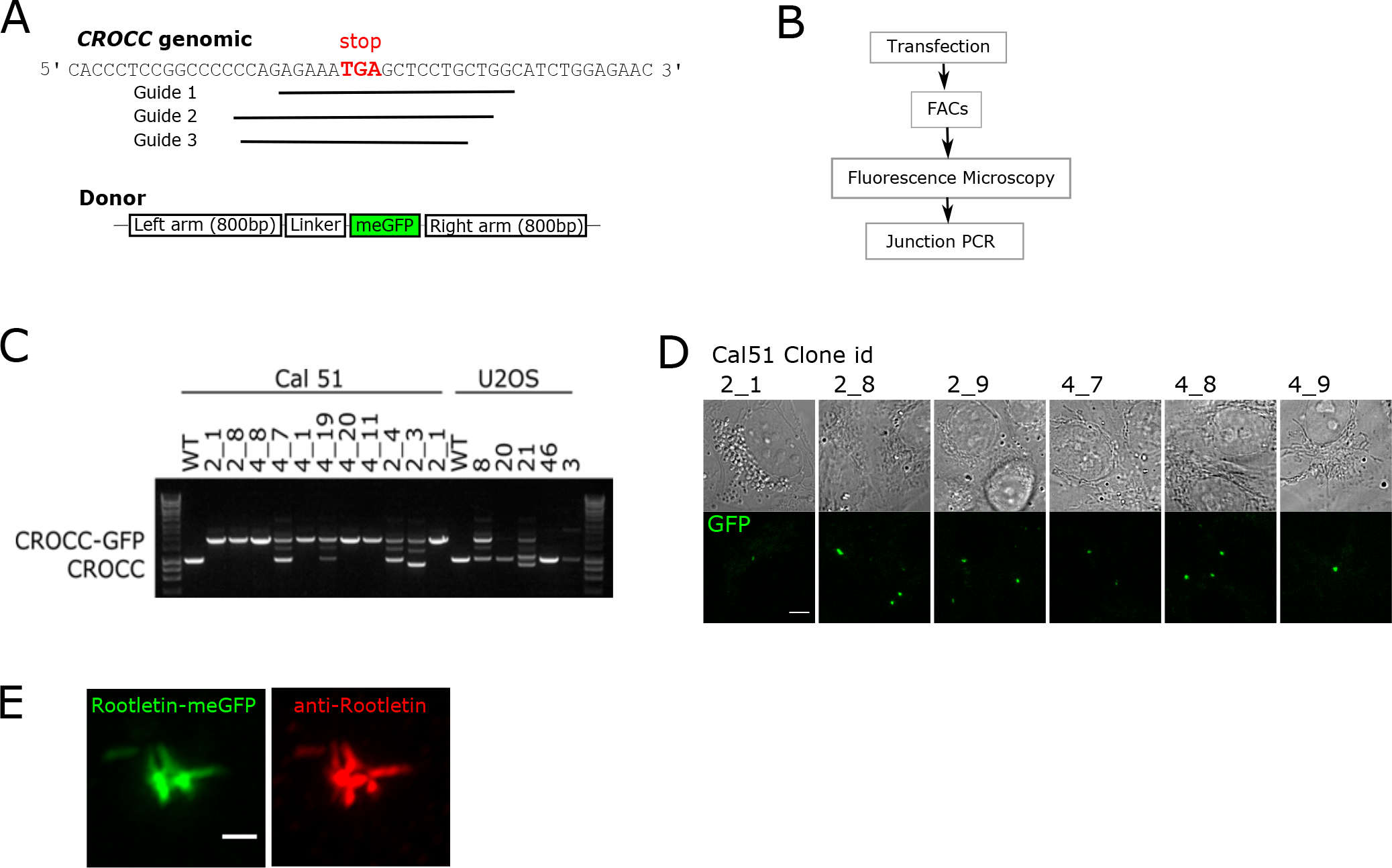
CRISPR Cas9 mediated tagging of endogenous rootletin / *CROCC* (related to Fig and Fig 3). (**A**) Schematic of guide RNAs targeting the STOP codon of *CROCC*, and a donor plasmid containing meGFP and homology arms. (**B**) Clones were screened sequentially by FACS sorting, fluorescence microscopy and junction PCR. (**C**) Example overlapping PCR screen of clones expressing rootletin-meGFP. Clone 4_1 was used in this study since it hashomozygous tagging of rootletin. Clones 4_7 and 20 are examples of heterozygous and negative clones respectively. (**D**) Representative fluorescence microscopy screening of clones expressing endogenous rootletin-meGFP. The bottom panel shows centrosomal fluorescence in positive clones. Scale bar 5μm. (**E**) Rootletin-meGFP centrosomal fluorescent signal closely resembles anti-rootletin antibody staining. The image shows clone 4_1 stained with anti-rootletin antibody (red) and imaged by Airyscan imaging. Scale bar 1μm.

## Author Contributions

R.M. conceived the project, performed the experiments, analysed results and wrote the manuscript.

## Acknowledgements

pEGFP Rootletin (Nigg pFL2(CW499)) was a gift from Erich Nigg (Addgene plasmid #41166). pX330-U6-Chimeric_BB-Cbh-hSpCas9 was a gift from Feng Zhang (Addgene plasmid #42230). pIRES GFP Centrin1 Hygro was a gift from Matthieu Piel (Addgene plasmid #64339). IMR-90 cells were a gift from Liam Cassidy. I thank the Cambridge Institute for Medical Research FACS facility for support. I thank Ashok R Venkitaraman and Paul M. W. French for guidance and feedback. Amy Emery, Petr Strnad, and Tom Miller provided critical comments on the manuscript. This work was funded by a Henry Wellcome Fellowship to R.M. (100090/12/Z).

## References

1. Schmidt KN, Kuhns S, Neuner A, Hub B, Zentgraf H, Pereira G. Cep164 mediates vesicular docking to the mother centriole during early steps of ciliogenesis. J Cell Biol. 2012;199: 1083–1101. doi:10.1083/jcb.201202126

2. Bahe S, Stierhof Y-D, Wilkinson CJ, Leiss F, Nigg EA. Rootletin forms centriole-associated filaments and functions in centrosome cohesion. J Cell Biol. 2005;171: 27–33. doi:10.1083/jcb.200504107

3. Bernhard W, De Harven E. [Electron microscopic study of the ultrastructure of centrioles in vertebra]. Z Zellforsch Mikrosk Anat Vienna Austria 1948. 1956;45: 378–398.

4. Bornens M, Paintrand M, Berges J, Marty MC, Karsenti E. Structural and chemical characterization of isolated centrosomes. Cell Motil Cytoskeleton. 1987;8: 238–249. doi:10.1002/cm.970080305

5. Graser S, Stierhof Y-D, Nigg EA. Cep68 and Cep215 (Cdk5rap2) are required for centrosome cohesion. J Cell Sci. 2007;120: 4321–4331. doi:10.1242/jcs.020248

6. Mayor T, Stierhof YD, Tanaka K, Fry AM, Nigg EA. The centrosomal protein C-Nap1 is required for cell cycle-regulated centrosome cohesion. J Cell Biol. 2000;151: 837–846.

7. Paintrand M, Moudjou M, Delacroix H, Bornens M. Centrosome organization and centriole architecture: their sensitivity to divalent cations. J Struct Biol. 1992;108: 107–128.

8. Chen JV, Kao L-R, Jana SC, Sivan-Loukianova E, Mendozça S, Cabrera OA, et al. Rootletin organizes the ciliary rootlet to achieve neuron sensory function in Drosophila. J Cell Biol. 2015;211: 435–453. doi:10.1083/jcb.201502032

9. Decarreau J, Wagenbach M, Lynch E, Halpern AR, Vaughan JC, Kollman J, et al. The tetrameric kinesin Kif25 suppresses pre-mitotic centrosome separation to establish proper spindle orientation. Nat Cell Biol. 2017;19: 384–390. doi:10.1038/ncb3486

10. Floriot S, Vesque C, Rodriguez S, Bourgain-Guglielmetti F, Karaiskou A, Gautier M, et al. C-Nap1 mutation affects centriole cohesion and is associated with a Seckel-like syndrome in cattle. Nat Commun. 2015;6: 6894. doi:10.1038/ncomms7894

11. Mazo G, Soplop N, Wang W-J, Uryu K, Tsou M-FB. Spatial Control of Primary Ciliogenesis by Subdistal Appendages Alters Sensation-Associated Properties of Cilia. Dev Cell. 2016;39: 424–437. doi:10.1016/j.devcel.2016.10.006

12. Panic M, Hata S, Neuner A, Schiebel E. The centrosomal linker and microtubules provide dual levels of spatial coordination of centrosomes. PLoS Genet. 2015;11: e1005243. doi: 10.1371/journal.pgen.1005243

13. Mahen R, Venkitaraman AR. Pattern formation in centrosome assembly. Curr Opin Cell Biol. 2012;24: 14–23. doi:10.1016/j.ceb.2011.12.012

14. Mahen R, Jeyasekharan AD, Barry NP, Venkitaraman AR. Continuous polo-like kinase 1 activity regulates diffusion to maintain centrosome self-organization during mitosis. Proc Natl Acad Sci U S A. 2011;108: 9310–9315. doi:10.1073/pnas.1101112108

15. Woodruff JB, Ferreira Gomes B, Widlund PO, Mahamid J, Honigmann A, Hyman AA. The Centrosome Is a Selective Condensate that Nucleates Microtubules by Concentrating Tubulin. Cell. 2017;169: 1066–1077.e10. doi:10.1016/j.cell.2017.05.028

16. Yang J, Liu X, Yue G, Adamian M, Bulgakov O, Li T. Rootletin, a novel coiled-coil protein, is a structural component of the ciliary rootlet. J Cell Biol. 2002;159: 431–440. doi:10.1083/jcb.200207153

17. Yang J, Gao J, Adamian M, Wen X-H, Pawlyk B, Zhang L, et al. The ciliary rootlet maintains long-term stability of sensory cilia. Mol Cell Biol. 2005;25: 4129–4137. doi:10.1128/MCB.25.10.4129-4137.2005

18. Yang J, Adamian M, Li T. Rootletin interacts with C-Nap1 and may function as a physical linker between the pair of centrioles/basal bodies in cells. Mol Biol Cell. 2006;17: 1033–1040. doi:10.1091/mbc.E05-10-0943

19. Fry AM, Meraldi P, Nigg EA. A centrosomal function for the human Nek2 protein kinase, a member of the NIMA family of cell cycle regulators. EMBO J. 1998;17: 470–481. doi:10.1093/emboj/17.2.470

20. Mardin BR, Lange C, Baxter JE, Hardy T, Scholz SR, Fry AM, et al. Components of the Hippo pathway cooperate with Nek2 kinase to regulate centrosome disjunction. Nat Cell Biol. 2010;12: 1166–1176. doi:10.1038/ncb2120

21. Sakaguchi H. PERICENTRIOLAR FILAMENTOUS BODIES. J Ultrastruct Res. 1965;12: 13–21.

22. Kuriyama R, Borisy GG. Centriole cycle in Chinese hamster ovary cells as determined by whole-mount electron microscopy. J Cell Biol. 1981;91: 814–821.

23. Sorokin SP. Reconstructions of centriole formation and ciliogenesis in mammalian lungs. J Cell Sci. 1968;3: 207–230.

24. Wang W-J, Soni RK, Uryu K, Tsou M-FB. The conversion of centrioles to centrosomes: essential coupling of duplication with segregation. J Cell Biol. 2011;193: 727–739. doi:10.1083/jcb.201101109

25. Lane HA, Nigg EA. Antibody microinjection reveals an essential role for human pololike kinase 1 (Plk1) in the functional maturation of mitotic centrosomes. J Cell Biol. 1996;135: 1701–1713.

26. Vassilev LT, Tovar C, Chen S, Knezevic D, Zhao X, Sun H, et al. Selective small-molecule inhibitor reveals critical mitotic functions of human CDK1. Proc Natl Acad Sci U S A. 2006;103: 10660–10665. doi:10.1073/pnas.0600447103

27. Mahen R, Koch B, Wachsmuth M, Politi AZ, Perez-Gonzalez A, Mergenthaler J, et al. Comparative assessment of fluorescent transgene methods for quantitative imaging in human cells. Mol Biol Cell. 2014;25: 3610–3618. doi:10.1091/mbc.E14-06-1091

28. Buendia B, Bré MH, Griffiths G, Karsenti E. Cytoskeletal control of centrioles movement during the establishment of polarity in Madin-Darby canine kidney cells. J Cell Biol. 1990;110: 1123–1135.

29. Mardin BR, Isokane M, Cosenza MR, Krämer A, Ellenberg J, Fry AM, et al. EGF-induced centrosome separation promotes mitotic progression and cell survival. Dev Cell. 2013;25: 229–240. doi:10.1016/j.devcel.2013.03.012

30. Piel M, Meyer P, Khodjakov A, Rieder CL, Bornens M. The respective contributions of the mother and daughter centrioles to centrosome activity and behavior in vertebrate cells. J Cell Biol. 2000;149: 317–330.

31. Schliwa M, Pryzwansky KB, Euteneuer U. Centrosome splitting in neutrophils: an unusual phenomenon related to cell activation and motility. Cell. 1982;31: 705–717.

32. Jean C, Tollon Y, Raynaud-Messina B, Wright M. The mammalian interphase centrosome: two independent units maintained together by the dynamics of the microtubule cytoskeleton. Eur J Cell Biol. 1999;78: 549–560. doi:10.1016/S0171-9335(99)80020-X

33. Au FKC, Jia Y, Jiang K, Grigoriev I, Hau BKT, Shen Y, et al. GAS2L1 Is a Centriole-Associated Protein Required for Centrosome Dynamics and Disjunction. Dev Cell. 2017;40: 81–94. doi:10.1016/j.devcel.2016.11.019

34. Graser S, Stierhof Y-D, Lavoie SB, Gassner OS, Lamla S, Le Clech M, et al. Cep164, a novel centriole appendage protein required for primary cilium formation. J Cell Biol. 2007;179: 321–330. doi:10.1083/jcb.200707181

35. Stinchcombe JC, Randzavola LO, Angus KL, Mantell JM, Verkade P, Griffiths GM. Mother Centriole Distal Appendages Mediate Centrosome Docking at the Immunological Synapse and Reveal Mechanistic Parallels with Ciliogenesis. Curr Biol CB. 2015;25: 3239–3244. doi:10.1016/j.cub.2015.10.028

36. Zwicker D, Decker M, Jaensch S, Hyman AA, Jülicher F. Centrosomes are autocatalytic droplets of pericentriolar material organized by centrioles. Proc Natl Acad Sci. 2014;111: E2636–E2645. doi:10.1073/pnas.1404855111

37. Euteneuer U, Schliwa M. Evidence for an involvement of actin in the positioning and motility of centrosomes. J Cell Biol. 1985;101: 96–103.

38. Wong M, Munro S. The specificity of vesicle traffic to the Golgi is encoded in the golgin coiled-coil proteins. Science. 2014;346: 1256898–1256898. doi:10.1126/science.1256898

39. Tinevez J-Y, Perry N, Schindelin J, Hoopes GM, Reynolds GD, Laplantine E, et al. TrackMate: An open and extensible platform for single-particle tracking. Methods San Diego Calif. 2017;115: 80–90. doi:10.1016/j.ymeth.2016.09.016

